# Ventral pallidal GABAergic neuron calcium activity encodes cue-driven reward-seeking and persists in the absence of reward delivery

**DOI:** 10.1101/2022.12.30.522319

**Authors:** Alexandra Scott, Dakota Palmer, Christelle A. Cayton, Iris Lin, Jocelyn M. Richard

## Abstract

Reward-seeking behavior is often initiated by environmental cues that signal reward availability. This is a necessary behavioral response; however, cue reactivity and reward-seeking behavior can become maladaptive. To better understand how cue elicited reward-seeking becomes maladaptive, it is important to understand the neural circuits involved in assigning appetitive value to rewarding cues and actions. Ventral pallidum (VP) neurons are known to contribute to cue elicited reward-seeking behavior and have heterogeneous responses in a discriminative stimulus (DS) task. The VP neuronal subtypes and output pathways that encode distinct aspects of the DS task remain unknown. Here, we used an intersectional viral approach with fiber photometry to record bulk calcium activity in VP GABAergic (VP GABA) neurons in male and female rats as they learned and performed the DS task. We found that VP GABA neurons are excited by reward-predictive cues but not neutral cues, and that this response develops over time. We also found that this cue-evoked response predicts reward-seeking behavior. Additionally, we found increased VP GABA calcium activity at the time of expected reward delivery, which occurred even on trials when reward was omitted. Together, these findings suggest that VP GABA neurons encode reward expectation and calcium activity in these neurons is predictive of the vigor of cue-elicited reward-seeking.

## INTRODUCTION

Environmental cues associated with rewards are powerful modulators of reward-seeking-behavior. Understanding how environmental cues can invigorate reward-seeking behavior is a necessary step in learning how cue reactivity, and subsequent reward-seeking, can become maladaptive. The ventral pallidum (VP) is a part of the ventral basal ganglia and an important node in the neural circuitry mediating cue-evoked reward-related behavior (Tindell et al., 2004, 2005; Smith et al., 2009; Richard et al., 2016; Ottenheimer et al., 2018; Richard et al., 2018; Ottenheimer et al., 2020a). VP neurons have heterogeneous responses to both rewards and cues predictive of reward. Most VP neurons are excited by reward-predictive cues, while a smaller population is inhibited by the same cues. Additionally, the firing rates of some cue-excited VP neurons can predict the vigor of reward-seeking in response to that cue (Richard et al., 2016, 2018). VP neurons are also sensitive to reward identity and prior reward history (Lederman et al., 2021). Whether the VP neurons encoding reward-predictive cues, reward-seeking vigor, and reward identity are the same or distinct subsets of VP neurons remains unknown. Therefore, exploring the neurochemical and projection-specific identity of VP neurons is an important step to understanding the broader neural circuitry that impacts reward-seeking behavior.

Recent research has begun to probe which VP neuronal cell-types contribute to distinct aspects of reward processing and reward-seeking behavior (Kupchik and Prasad, 2021). The VP contains 3 major neuronal subtypes: cholinergic, glutamatergic, and gamma-aminobutyric acidergic (GABA) (Root et al., 2015). VP glutamatergic neurons constrain reward-seeking and promote aversion related behavior (Faget et al., 2018; Tooley et al., 2018; Stephenson-Jones et al., 2020). In contrast, VP GABA neurons are excited by rewards and Pavlovian reward-paired cues (Heinsbroek et al., 2020; Stephenson-Jones et al., 2020). Activation of VP GABA neurons drives positive reinforcement as well as cued drug-seeking and relapse (Faget et al., 2018; Heinsbroek et al., 2020; Prasad et al., 2020; Farrell et al., 2022). Inhibition of VP GABA neurons can induce place avoidance and dampen risky decision making (Faget et al., 2018; Heinsbroek et al., 2020; Farrell et al., 2021). Yet, how VP GABA neurons develop responses to reward-predictive cues over time, or if their activity predicts the vigor of reward-seeking, remains unknown.

Here we examined whether the calcium activity of VP GABA neurons encodes cues predictive of reward availability and how this activity is related to subsequent reward-seeking behavior. Rats were trained in a discriminative stimulus (DS) task with liquid sucrose reward while we recorded population-level calcium activity of VP GABA neurons using fiber photometry. We found that VP GABA neurons develop a calcium response to reward-predictive cues as rats learn the DS task. VP GABA calcium activity was found to respond following the reward-predictive cue, following the action required to obtain reward, and during reward consumption. Calcium increases post-cue and operant action persisted even on trials where reward was omitted. Finally, VP GABA cue-evoked calcium activity was shown to correlate with behavioral responses that measure the vigor of reward-seeking. These results suggest that VP GABA neurons encode environmental cues predictive of sucrose reward and this cued calcium response predicts the vigor of reward-seeking behavior.

## MATERIALS AND METHODS

### Subjects

Male and female Long Evans rats (n=21, 9 male, 12 female; Envigo), weighing 250–275 grams at arrival, were individually housed in a temperature- and humidity-controlled colony room on a 12-hr. light/dark cycle. Rats were food restricted starting three days prior to the initiation of training in the DS task and until experiments were done. Rats received 5% of their body weight in food each day and the amount of food was adjusted daily to maintain rats at ~90% of their free-feeding body weight. All experimental procedures were approved by the Institutional Animal Care and Use Committee at the University of Minnesota and were carried out in accordance with the guidelines on animal care and use of the National Institutes of Health of the United States.

### Surgeries

During surgery, rats were anesthetized with isoflurane (5%) and placed in a stereotaxic apparatus, after which surgical anesthesia was maintained with isoflurane (0.5–2.0%). Rats received preoperative injections of carprofen (5 mg/kg) for analgesia and cefazolin (75 mg/kg) to prevent infection. To achieve cell-type specific calcium indicator expression we used a dual viral approach combining a glutamate decarboxylase 1 (GAD1) promoter virus for the expression of Cre recombinase with a virus for expression of GCaMP6f in a cre-dependent manner (Liu et al., 2013; Wakabayashi et al., 2019; Shields et al., 2021). Rats (n=19, 9 male, 10 female) received 0.8 uL of a 1:1 mixture of AAV9-Syn-Flex-GCaMP6f (2.1 * 10^13 GC/mL; Addgene) and AAV8-GAD1-cre (8.29×10^13 GC/mL; University of Minnesota Viral Vector Core) was injected unilaterally into VP. Syringes for viral delivery and optical fiber implants were aimed at the VP using the following coordinates relative to bregma: +0.3 mm AP, +/− 2.3 mm ML, −8.3 mm DV. A subset of these rats (n=4, 2 male, 2 female) also received, contralateral to the primary recording location, an infusion of AAV9-Syn-Flex-GCaMP6f and fiber implant into VP and a retrograde AAV8-GAD1-cre (AVVrg-GAD1-Cre;1.03×10^14 GC/mL; University of Minnesota Viral Vector Core) into LH (−2.9 mm AP, 1.8 mm ML, −8.4 mm DV). This combination of injections did not lead to GCaMP6f expression. Thus, these recordings are included in the “no expression” control data. For external controls (n=2, 2 female) 0.8 uL of a 1:1 mixture of EGFP (AAV9- Hsyn-DIO-EGFP;4.6*10^13 GC/mL; Addgene) and AAV8-GAD1-cre was injected unilaterally into VP. Virus was delivered through 28-gauge injectors at a rate of 0.1 μl per min. Injectors were left in place for 10 min following the infusion to allow virus to diffuse away from the infusion site. Following viral injections, rats used for fiber photometry measurements and behavioral testing (n=15; 6 males and 9 females) received fiber implants (.48 NA, 400 micron, 9mm; Doric) in VP for optical measurement of calcium activity. Implants were secured to the skull with bone screws and dental acrylic. Rats recovered for at least one week before beginning handling. To allow for sufficient viral expression, rats recovered for at least 4 weeks before behavioral training or brain extraction for RNAscope.

### Discriminative Stimulus Task

Four to six weeks after surgery, rats were trained in a discriminative stimulus (DS) task (Ottenheimer et al., 2018; Richard et al., 2018; Gómez-A et al., 2022). The DS task is an instrumental task where rats learn to discriminate between reward-predictive (DS) and neutral cues (NS). Auditory cues consisted of a siren or white noise, with cue assignments counterbalanced across subjects. Rats received reward if they entered the port during the DS cue but not the NS cue. Over time rats learn to discriminate between the cues, responding almost exclusively to the DS.

#### Initial Training (Port Training and Stages 1-4)

Prior to training with cues, rats underwent port training in the operant boxes while tethered to the fiber optic patch cables to acclimate them to both the cables and operant boxes. In port training sessions, animals received 30 trials of 10% sucrose delivery. For each trial, once sucrose reward was retrieved from the port, another trial was initiated. Once rats met the criteria of 30 reward retrievals in 1 hour, they transitioned to Stage 1 of DS task training. In Stages 1-4 of the DS task, each session consisted of 30 DS trials. Each trial consisted of a window in which an auditory cue (the DS; siren or white noise) signaled 10% sucrose availability for up to 60 seconds (Stage 1), 30 seconds (Stage 2), 20 seconds (Stage 3), or 10 seconds (Stage 4). The DS cue played until the rat entered the port or the availability window was over. Inter-trial intervals (ITI) were pseudo-random and adjusted so that the average time between the start of each DS presentation was 120 seconds (i.e. as the DS length decreased, the ITI lengthened from 60-110 seconds). Rats were required to respond during at least 70% of DS presentations (0.70 response probability) to pass each stage. After meeting these criteria for Stage 4, rats moved onto the full version of the DS task.

#### Full DS Task Training (Stage 5)

The full version of the DS task (Stage 5) consisted of 60 trials, 30 of which were DS and 30 of which were NS. The DS was presented for a maximum of 10 seconds, or until the animal entered the port. The NS played for 10 seconds independent of animal behavior. DS or NS trial order was determined pseudo-randomly. Inter-trial intervals were determined pseudo-randomly with an average interval time of 50 seconds (30-70 seconds). A criterion of 0.70 DS response probability and 0.30 NS response probability was required for animals to complete Stage 5.

#### Probe Trials and Extinction

Following completion of Stage 5 testing, we added in a 1 second delay between the port entry and reward delivery, as well as probe trials for 50% of DS presentations. During probe trials, port entries during the DS were not followed by reward delivery. This allowed us to better determine whether we saw distinct calcium responses to the cue presentation, port entry, and initial lick bouts. Rats underwent 2 days of testing under these conditions. Finally, animals underwent 3 days of extinction in which sucrose was no longer delivered.

### *In vivo* Calcium Recordings

Fiber photometry recordings were done throughout training in the DS task. Recordings were conducted according to methods described previously (Saunders et al., 2018). Rats were first acclimated to the tethering cable (0.48, 400-micron core optical fiber) and the operant box. During recording sessions, the tethering optical fiber is connected to a fluorescent mini cube (Doric Lenses, Canada) which transmits excitation light from both a 465 nm LED and a 405 nm LED. 465 nm excitation stimulates calcium-dependent GCaMP6f fluorescence, and 405 nm excitation stimulates GCaMP6f fluorescence in a calcium-independent fashion. GCaMP6f fluorescence travels to the mini cube via the tethering cable, passes through a GFP emission filter and is amplified and focused onto a high sensitivity photodetector. A real-time signal processor (RZ5P, Tucker-Davis Technologies) running Synapse software was used to modulate the output of each LED and record photometry signals. Before each session, the power level for each LED was measured and adjusted to a range of approximately 10-25 microwatts (Meng et al., 2018). The offset for each animal was kept constant, between 20-30 mA (Tucker-Davis Technologies user manual), and the level was adjusted to obtain the appropriate power. The 465 nm LED driver frequency was set to 211 Hz while the 405 nm LED driver frequency was set to 531 Hz. This allowed for demodulation of the separate LED signals, yielding a calcium-dependent signal and a calcium-independent signal (isosbestic signal; (Tian et al., 2009; Dana et al., 2018; Patel et al., 2019) to use as an internal control for each animal. Task events (cue presentations, port entries and lick bouts) were time stamped in the photometry data file via a TTL signal generated by the behavioral equipment (Med Associates Super Port and TTL panel). Video recordings were simultaneously collected from each operant box (Amcrest).

Calcium-dependent fluorescent changes were detected in the calcium dependent channel (465nm), whereas calcium-independent fluorescent changes were detected in both the dependent and independent (405 nm, isosbestic) calcium channels. VP GABA bulk activity is represented as a change in fluorescence over time. Calcium dependent and independent channels were fit and combined to control for any artifacts not associated with neuronal activity. This combined signal (delta F/F) was normalized in peri-event time windows for each trial, where baseline activity 5 seconds before cue presentation was used to calculate a z-score for each delta F/F data point in the peri-event window.

### Behavioral Analysis

DS and NS response probabilities were calculated for each cue type as the ratio between the number of trials where the animal entered the port within 10 seconds of cue onset to the number of overall trials, for each specific cue type. We used port entry latency as a behavioral proxy for reward-seeking vigor and motivation (Richard et al., 2018; Lederman et al., 2021). Port entry latency was calculated as the difference between port entry time points and cue onset time points. Port entry latency was only calculated on trials where animals entered the port within 10 seconds of cue presentation, allowing for a more consistent measure of latency across stages. One animal was excluded for lack of cue discrimination after 12 days of Stage 5 training. To examine how the port entry probability developed across training, two linear mixed-effects models were run. For training sessions between Stages 1-4, the model was fit with fixed effects for day and sex, and a random effect of subject. For Stage 5 sessions, the model was fit with fixed effects for day, sex, and cue type, and a random effect of subject. A paired t-test was run for criteria day port entry probability and latency, between cue types.

### Fiber Photometry Analysis

#### Post-cue response across training

Fluorescent values and behavioral timestamps were exported to MATLAB. Each training session was analyzed in a custom MATLAB script (available upon request). Signals driven by 465 nm and 405 nm LEDs were down sampled to 40 Hz, fit to one another, and combined (465 nm - fitted 405 nm/fitted 405 nm; delta F/F) to control for any artifacts not associated with neuronal activity. Z-scores of the delta F/F photometry signals for each trial were calculated based on the mean and standard deviation of fluorescent values 5 seconds before the cue. We used an area under the curve analysis on z-scored trial traces to collect a single value between 0- and 5-seconds post-cue (when the bulk of the DS calcium event occurred) to run linear mixed effects models. To examine how the calcium signal developed across training, two linear mixed effects models were run. For training sessions between Stages 1-4, the model was fit with fixed effects for day and sex, and a random effect of subject. For Stage 5 sessions, the model was fit with fixed effects for day, sex, and cue type and random effect of subject. Depending on the best-fitting covariance model (Verbeke, 1997), the degrees of freedom may be a non-integer value. Any significant interactions were further assessed with pairwise comparisons with Sidak corrections for multiple comparisons.

#### Isolating post-cue responses to behavioral events

To isolate calcium signals associated with the cue from that of the port entry and reward delivery, we ran an encoding model on Z-scored calcium peri-event time windows (Parker et al., 2016). The analysis isolates each timestamped event (cue presentation, port-entry, initial lick onset) and relates that to each sampling point recorded from the calcium signal. To do this, event onset timestamps were transformed into binary indicators for each sampling point. Then, linear regression was run using the z-scored, stage 5, GCaMP6f recordings as the response variable and the transformation of the timestamps of the cue presentation, port entry, and initial licks as the predictor variables. This regression model output provided response kernels for each distinct event. AUC values were then acquired from regression kernels and student’s t-tests were run defining the AUC’s significance from null.

#### Post-cue correlation to reward-seeking motivation

To determine if DS-evoked neuronal activity predicts the latency of reward-seeking, our behavioral proxy of reward-seeking motivation, we ran a linear correlation analysis relating the peri-DS Z-scored delta F/F calcium signal from Stage 5 trials to trial-by-trial port entry latency (Pearson’s correlation coefficients) for each rat. Because the encoding model described above revealed significant port-entry related increases in calcium activity after, but not before, port entry, for our correlation analysis we excluded fluorescence recorded from any post-port entry time points. We also shuffled these data to create random control data for comparison to the true calcium and latency relations. Comparison of the shuffled and unshuffled correlations were done in 0.10 second time bins with multiple t-tests with Sidak corrections for multiple comparisons.

### Histology

#### Validation of fiber placement and region-specific expression

Following completion of behavioral testing with fiber photometry recordings, animals (n=15, 6 males and 9 females) were deeply anesthetized with pentobarbital and were perfused intracardially with 0.9% saline followed by 4% paraformaldehyde. Brains were removed, post-fixed in 4% paraformaldehyde for 4–24 hr., cryoprotected in 30% sucrose for >48 hours, and sectioned at 40 um on a microtome. Sections were then washed with PBS, wet-mounted on coated glass slides in PBS, air-dried, cover slipped with Vectashield mounting medium with DAPI, and imaged on a fluorescent microscope. We verified the location of injection or optical fiber recording sites using anatomical markers of VP (anterior commissure, lateral ventricle size). Additionally, fiber implants in the contralateral VP were originally implanted in a subset of subjects to record from lateral hypothalamus projecting VP GABA neurons, however, viral expression was not obtained, and no fluorescent protein was expressed in the cells. We used recordings from these animals (n=4, 2 male, 2 female) as “no expression” control sessions. Placement of fiber was assessed for these control recordings as well and confirmed to fall within VP.

#### Validation of cell-type specific GCaMP6f expression

Animals designated for RNAscope analysis (n=6, 3 females, 3 males) were deeply anesthetized with pentobarbital and decapitated. Fresh tissue was extracted and flash frozen in dry-ice-cooled isopentane. Brains were stored in a −80-degree Celsius freezer for up to 1 year. Tissue was stored until sectioning of tissue was done on the cryostat (coronal sections, 16 um) and jump mounted for RNAScope processing. Tissue was processed to determine the location and extent of viral expression and neural identity of the neurons recorded. To validate our viral approach, we used RNAscope^®^ *in situ* hybridization with probes for *Gad1, Slc17a6 (Vglut2*) and *GCaMP6* (Fig. 1D-E). The channels were respectively: GCaMP6 probe (488 nm excitation), glutamatergic probe (Slc17a6, 555 nm excitation), GABAergic probe (Gad1, 647 nm excitation). Imaging of the tissue was done on a confocal (Nikon upright c2, UIC-UMN) at 20x with 2x digital zoom, and settings were optimized with the help of the imaging core. The exact same acquisition settings were used on all tissue samples. HALO software (Indica Labs) quantified the amount of cell specificity achieved from viral injections. Briefly, total cell number and location within an image was determined by nuclear staining (DAPI). Probes (Gad1, vGlut, GCaMP6) are assigned their respective fluorophores within the software and the software compares the location of each fluorophore to the location of the DAPI labeled cells to confirm fluorophore expression is associated with a self-defined nuclei and cell size. HALO analysis software and software settings were optimized for the tissue from all animals and then settings were used to batch analyze cell counts of GCaMP6f positive cells. No difference was seen in expression between different subregions of VP. The percentage and number of each defined cell identity of interest were determined for 3 sections per animal to determine cell specificity within and between all subjects.

**Figure 1.**
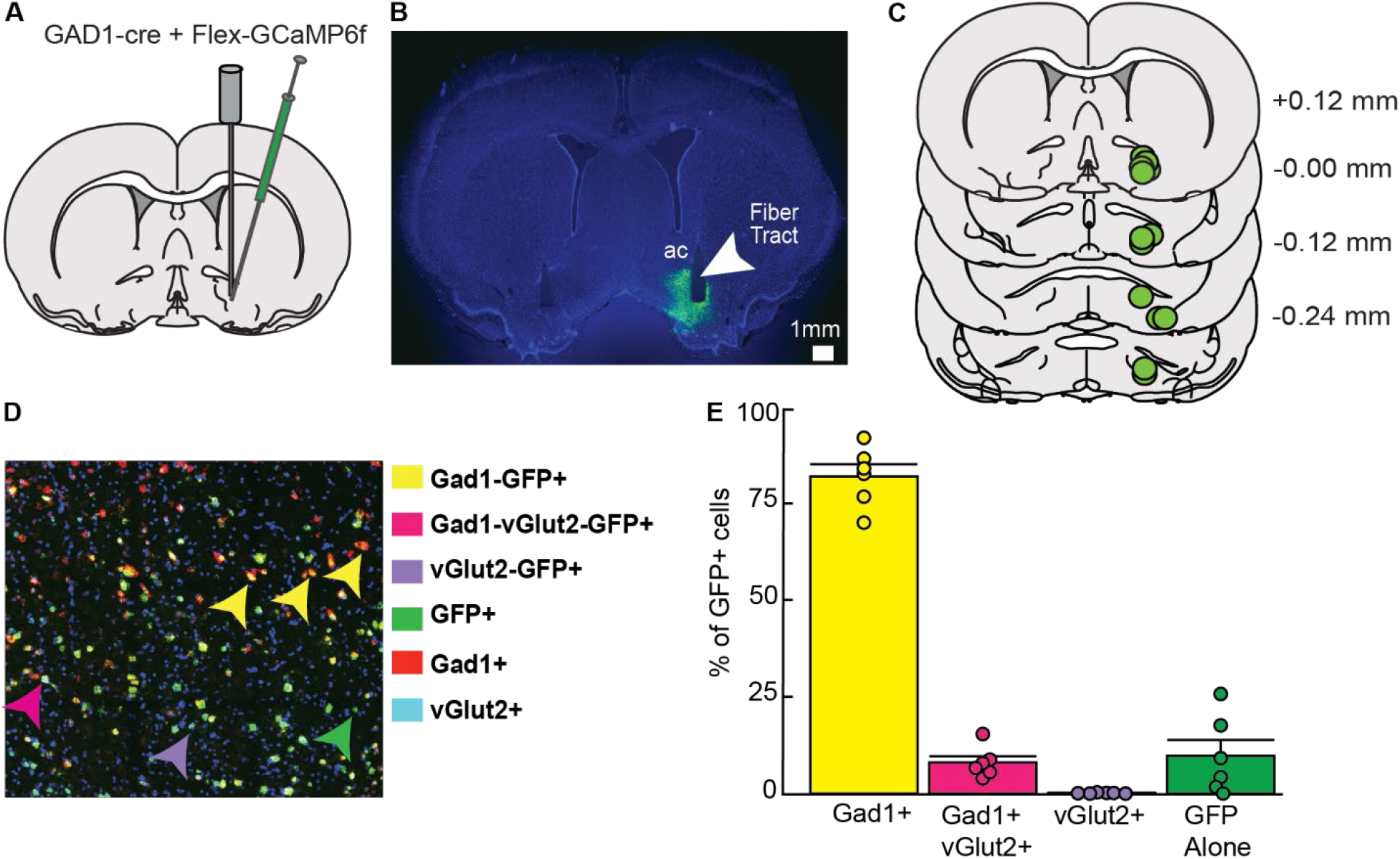
Anatomical and cellular characterization of GCaMP6f expression in ventral pallidum (VP). (A) Diagram of intersectional viral approach and fiber optic implant used for cell type-specific calcium recordings in the VP. (B) Representative image of virally expressed GCaMP6f and optical fiber tract placement in VP for calcium activity recordings (right side), and for no-expression control recordings (left side). (C) Verification of fiber and expression at calcium activity recording sites in VP-containing coronal sections. Values listed indicated distance (mm) from bregma. (D) Representative image of RNAscope processed tissue with arrows denoting location of each defined GCaMP6f expressing (GFP+) cell type. (E) Quantification of the % GFP+ cells expressing Gad1, vGlut2 or both cell-type specific markers, n=6; 3 VP sections utilized for each subject’s average (82.09% +/− 3.14% Gad1, 0.12% +/− 0.05% of glutamatergic, 8.03% +/− 1.63% Gad1/Vglut2/GFP, 9.75 % +/− 4.12% of just GFP+. Data are mean +/− SEM. Dots represent individual rats.

## RESULTS

### Viral mixture and fiber implantation specifically targets ventral pallidal GABAergic neurons

A mixture of DIO-GCaMP6f and GAD1-Cre viruses were infused into VP to express the genetically encoded calcium indicator (GCaMP6f) specifically in VP GABA neurons. For measurement of calcium-related fluorescence, an optical fiber was implanted into VP (Fig. 1A–1B). All rats that underwent calcium activity recording had viral expression and fiber placements within VP (Fig. 1C). To validate the cell-type specificity of our viral approach, we used RNAScope *in situ* hybridization with probes for *Gad1, Slc17a6 (Vglut2*) and GCaMP in rats that did not receive fiber implants (Fig. 1D). Because no difference was seen in expression or selectively in different subregions of VP, these data were pooled for analysis. Overall, GCaMP6f (GFP +) and Gad1 positive cells made up 90.12 % of GFP + cells (82.09% +/− 3.14% Gad1/GFP, 8.03% +/− 1.63% Gad1/Vglut2/GFP; Fig. 1E). Therefore, the cells we were recording with fiber photometry during the DS task were primarily and selectively GABAergic. In contrast, only 0.12% +/− 0.05% of GFP + cells were exclusively glutamatergic. Finally, 9.75 % +/− 4.12% of cells were GFP+ with no glutamatergic or GABAergic probe expression.

### Rats discriminate between reward-predictive and neutral cues

Rats (n=12; 6M, 6F) were trained on an operant discriminative stimulus task where reward availability was signaled by a reward-predictive cue (DS). Rats received a 10% sucrose reward if they entered the operant chamber port (port entry) during DS presentations, but not if they entered during the control cue (NS) or non-cue periods (Fig. 2A). During Stages 1-4 (DS alone) of the task, rats received presentations of just the DS, which decreased in length over sessions from 60 to 10 seconds incrementally. As expected, port entry probability during the DS increased significantly across sessions during this phase of training (Fig. 2B; session effect: F(1,98.36) = 93.01, p < 0.001).

**Figure 2.**
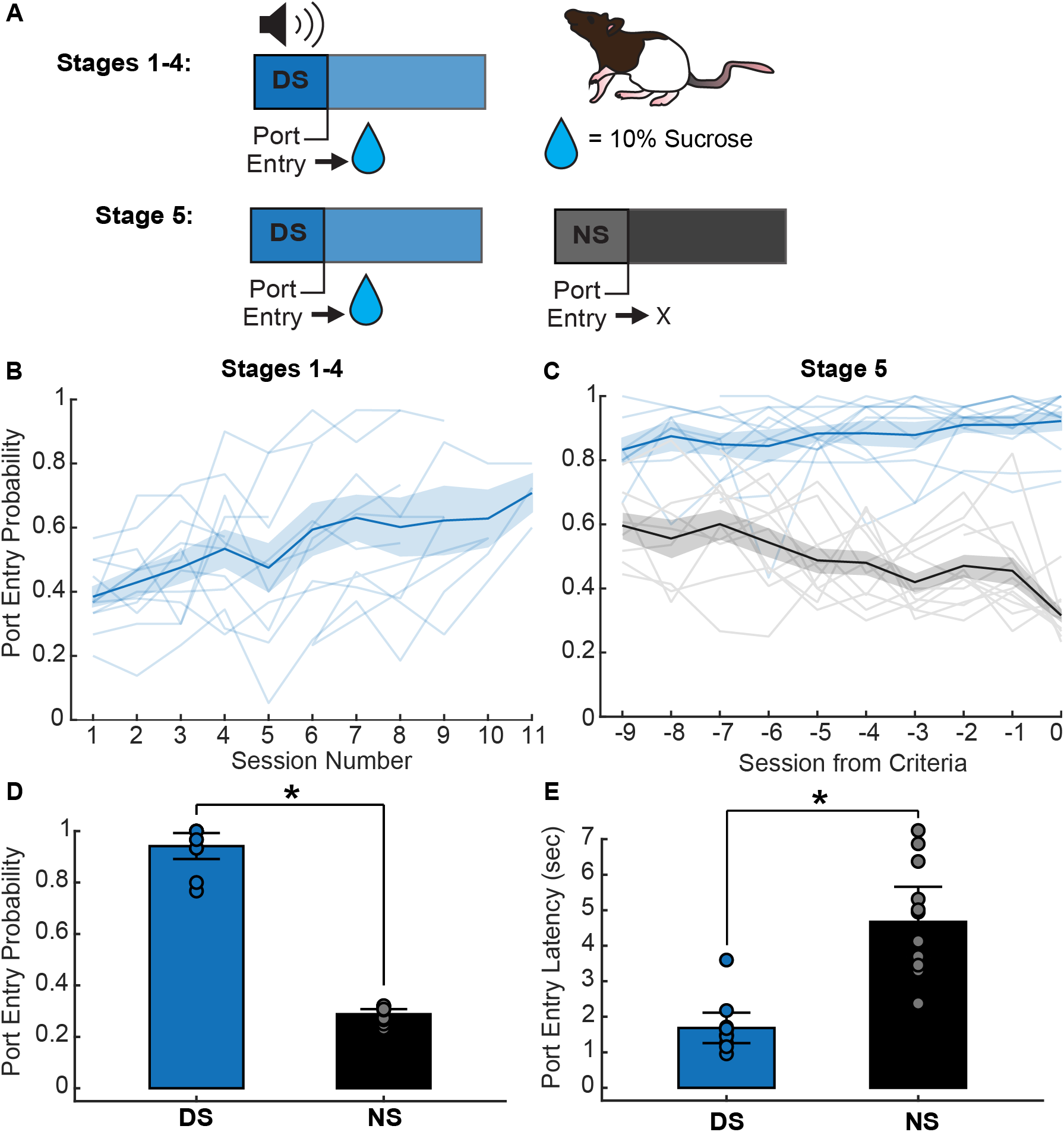
Behavioral task and training data. (A) DS behavioral task trial design using 10% sucrose reward. (B) DS response probability (blue) during stages 1-4 of DS task (session effect: F(1,98.36) = 93.01, p < 0.001). The line with shading indicates mean +/− SEM. Lines alone indicate individual subjects. (C) Average DS (blue) and NS (black) response probability across the last 10 days of stage 5 training (9 days before criteria is reached; session*cue effect: F(1,218.46) = 25.32, p < 0.001). The line with shading indicates mean +/− SEM. Lines alone indicate individual subjects. (D) Average DS and NS response probability on the day discrimination criteria was reached (paired t-test, *p < 0.001). Data are mean +/− SEM. Dots represent individual rats. (E) Average DS and NS port entry latency on day discrimination criteria reached (paired t-test, *p< 0.001). Data are mean +/− SEM. Dots represent individual rats.

Once rats entered the port during at least 70% of DS presentations in Stage 4, they progressed to the full version of the task (Stage 5), in which the NS control cue was introduced (Fig. 2A). Rats were deemed capable of discriminating between the DS and NS when they met criteria of a minimum of 70% percent DS response probability and a maximum of 30 % NS response probability. One animal was excluded for lack of cue discrimination after 12 days of stage 5 training.

As expected, rats learned to discriminate between the DS and NS cues over time (Fig. 2C; interaction of session and cue, F(1,218.46) = 25.32, p < 0.001), increasing their probability of entering the port during the DS, and decreasing their probability of entering the port during the NS. In addition to responding more frequently to the DS than the NS when they met criteria (Fig. 2D; p < 0.001), rats also entered the port more quickly following the DS compared to the NS on trials when they made a port entry (Fig. 2E; p < 0.001).

### Ventral pallidal GABA neurons develop a calcium response following the onset of the reward-predictive cue but not the neutral cue

Fiber photometry measurements of calcium activity were conducted throughout training to assess the development of responses across learning of the DS task. Using these methods, as rats learned the task, we observed VP GABA neurons develop a calcium response following the onset of the DS (Fig. 3A) but not the NS (Fig. 3B). The VP GABA neuronal calcium response is not apparent in response to the DS during the first session of training (Fig. 3C). Area under the curve (AUC) was calculated for each individual rat’s average Z-scored calcium trace during each session of training. For initial training with just the DS, we observed a significant effect of training session on the average AUC prior to the introduction of the NS (Fig. 3F; main effect of session, F(8,63.53) =5.16, p < 0.001). By the first day of Stage 5, when the NS is introduced, an increase in calcium activity occurs soon after DS onset and persists for nearly 10 seconds (Fig. 3D, blue line). Cue-evoked calcium activity in response to the NS is reduced relative to the DS (Fig. 3D, black line). On the day subjects met behavioral criteria for cue discrimination (70% or greater response to the DS, 30% or less for the NS) the VP GABA calcium response post-DS is robust and persists for closer to 5 seconds, whereas the post-NS response is minimal in amplitude and duration. AUC analysis for the last 10 sessions of Stage 5 training shows a significant difference between DS and NS calcium responses (Fig. 3G; main effect of cue, F (1,194.18) =46.64, p < 0.001). No effect of cue was seen in control recordings (GFP and no-expression controls; main effect of cue, F(1,48.37) =0.18, p=.68). These findings suggest VP GABA neurons develop a response to reward-predictive cues while learning the DS task, but do not respond to neutral cues never paired with reward.

**Figure 3.**
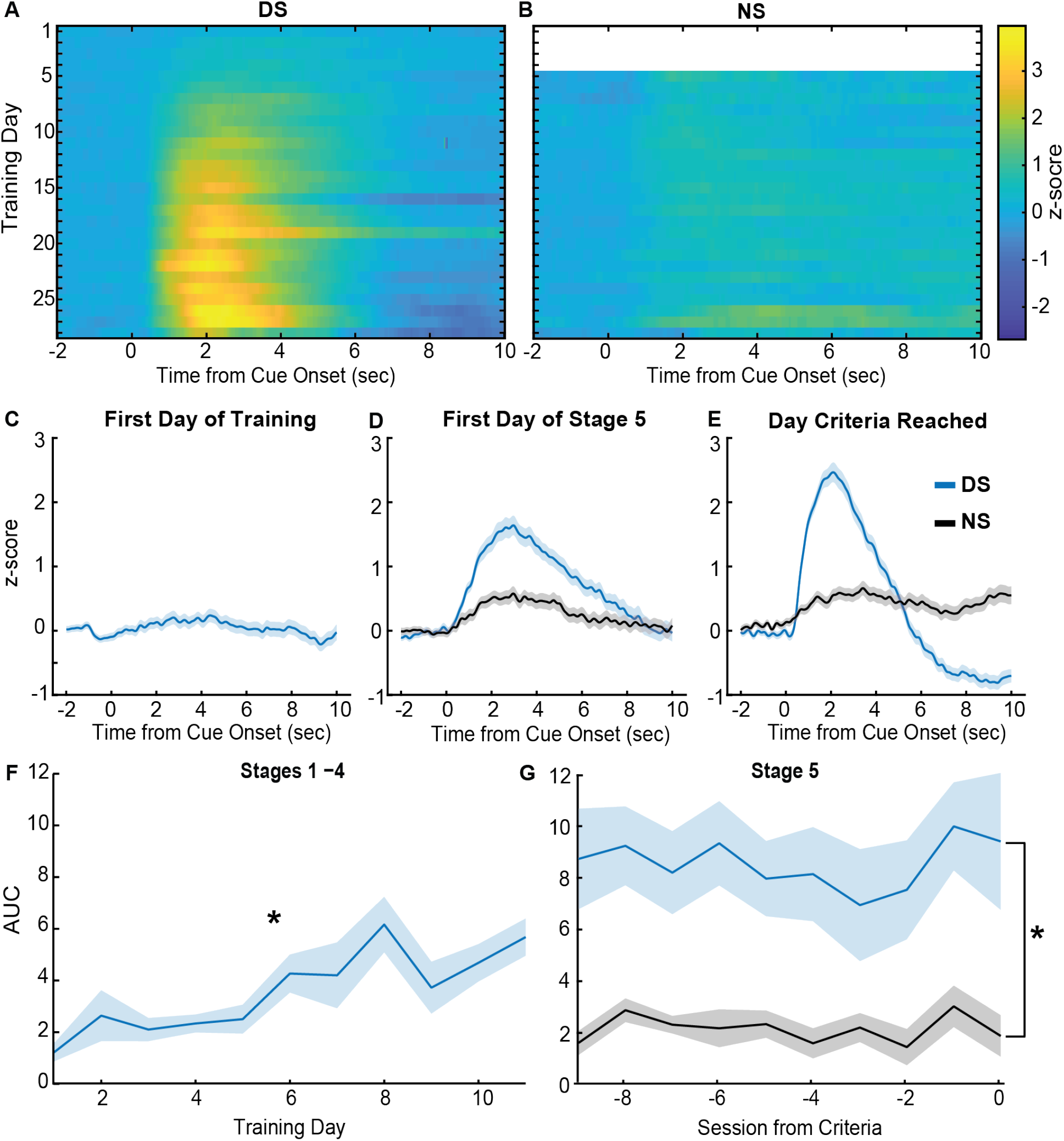
Calcium activity during training. (A) Heatmap of average Z-scored delta F/F response from VP GCaMP6f during the peri-DS cue time window (x-axis) across training days (y-axis) of all stages (n=12 rats) (B) Heatmap of z-scored delta F/F response during the peri-NS cue time window across all Stage 5 training days (n=12 rats). (C) Average peri-DS calcium trace on day 1 of training (n=12 rats; +/− SEM). (D) Average peri-cue (DS=blue, NS=black) calcium trace on the first day of Stage 5 (n=12 rats; +/−SEM). (E) Average peri-cue calcium trace on the day each animal met criteria (n=12; +/− SEM). (F) Average AUC (0-5 seconds) of post-DS calcium traces across animals (n=12; +/− SEM) during Stages 1-4 (main effect of session, effect: F(8,63.53)=5.16, *p < 0.001). (G) Average AUC (0-5 seconds) of post-cue calcium traces for the DS (blue) and NS (black) across animals (n=12; +/− SEM), during the last 10 days of Stage 5 training (main effect of cue, F(1,194.18)=46.6437, *p < 0.001).

### VP GABA neuronal response post reward-predictive cue is associated with cue onset, operant behavior, and initial reward consumption

Prior work from electrophysiological recordings in VP has found that DS-excited single units increase their firing at a median time of 90 milliseconds after cue onset and return to baseline within 1 second, even for units with the longest duration responses (Richard et al., 2016, 2018). In contrast, we observed changes in fluorescence from population-level calcium activity in VP GABA neurons, starting approximately 1 second after cue onset and persisting in the average traces for multiple seconds. While GCaMP6f is a fast calcium indicator (Chen et al., 2013), in terms of both rise and decay kinetics, it still introduces slower dynamics compared to electrophysiological recordings (Wei et al., 2020). Prior work from electrophysiological recordings in the DS task has also found that VP single units are responsive to reward-seeking actions and at the time of reward consumption (Richard et al., 2016, 2018) and may encode the relative value of primary rewards or reward prediction errors (Ottenheimer et al., 2018, 2020a, 2020b) in addition to the incentive motivational value of cues. Therefore, it is likely that the calcium activity of VP GABA neurons is sensitive to multiple behavioral epochs following cue onset. That is, multiple calcium responses may contribute to the prolonged calcium trace seen when trials are averaged. When we sorted individual trials by the animals’ latency to enter the port, we observed both increased fluorescence that was time-locked to the cue, as well as an increase that appeared to be time locked to the port entry after the DS presentation (Fig. 4A). We observed a similar pattern when we plotted the trials time-locked to port entry (PE) and sorted by port entry latency (Fig. 4B). Moreover, when trials that were time-locked to initial lick responses were sorted by port entry latency, we observed a distinct calcium response that occurs after initial lick onset, at least on trials where latency between reward delivery (time of PE) and licking behavior is longer (Fig. 4C). Therefore, the extended calcium responses we see following DS onset could be related to all three behavioral epochs: sensing the cue, the operant action, and the initial reward consumption.

**Figure 4.**
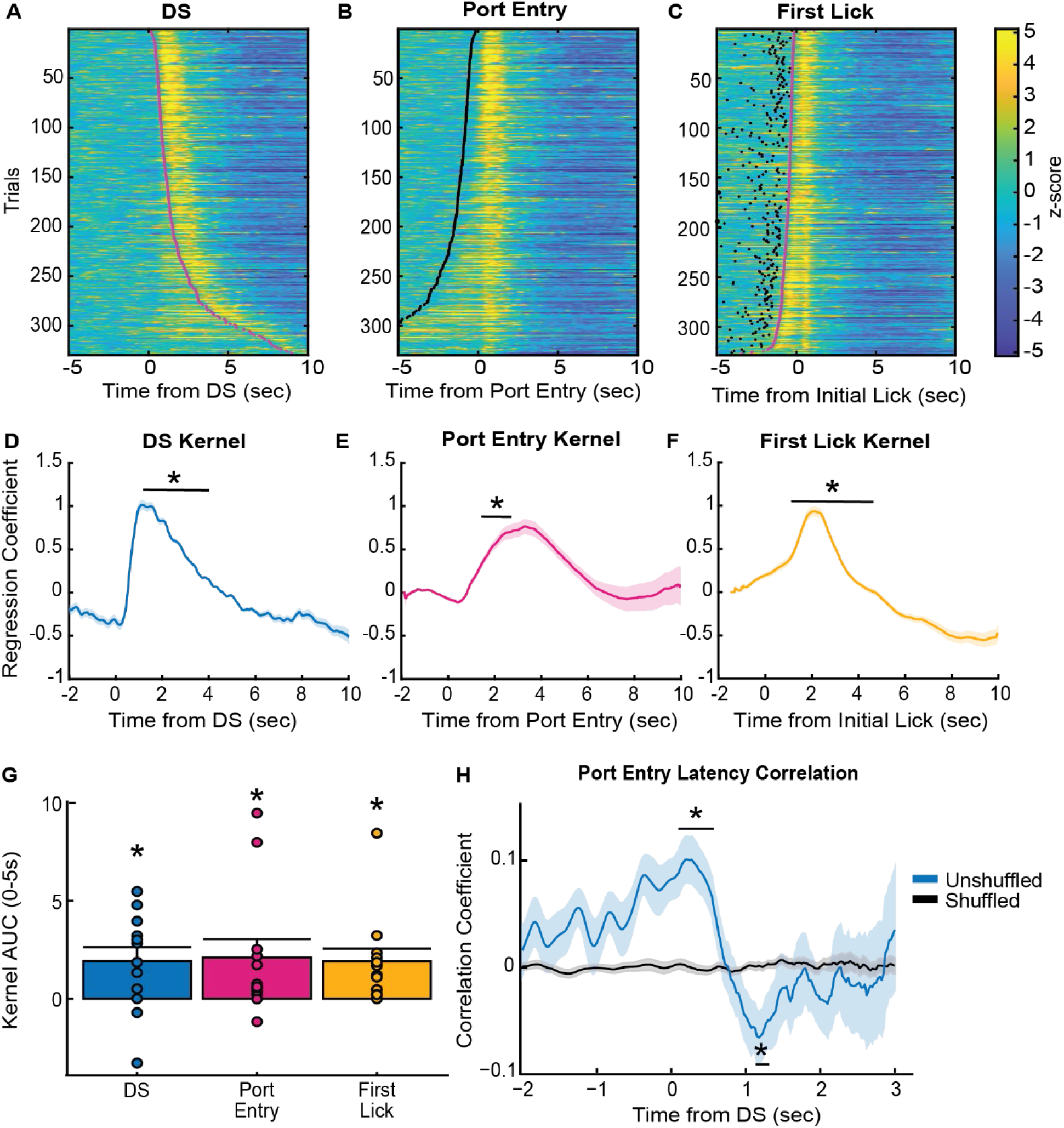
Event-related responses during performance of the DS task. (A) Heatmap of Z-scored responses from Stage 5 of a representative animal, trials sorted by PE latency, time-locked to the DS cue onset. Magenta dots represent the time of port entry. (B) Heatmap of Z-scored responses from Stage 5 of a representative animal, trials sorted by PE latency, time-locked to port entry. Black dots represent cue onset. C) Heatmap of Z-scored responses from Stage 5 of a representative animal, sorted by PE latency, time-locked to initial lick onset. (D) Average DS cue onset kernel of lasso regression coefficients across all animals (n=12), significant difference from zero between 0.90-3.40 sec time bins (*ps<0.05) (E) Average port entry onset kernel of lasso regression coefficients across all animals (n=12), significant difference from zero between 1.40-2.40 sec time bins (*ps<0.05) (F) Average initial lick onset kernel of lasso regression coefficients across all animals (n=12), significant difference from zero between 1.30-3.60 second time bins (*ps<0.05). (G) Average AUC of each regression kernel across animals (n=12), t-test comparing each AUC to zero. Mean + SEM, dots indicate individual subjects. (H) Linear Pearson’s correlation coefficients for animal’s trial latency to DS evoked VP GABA calcium activity correlation (unshuffled= blue, shuffled=black). Time bin analysis indicated that true (unshuffled) coefficients were significantly different from zero between 0.00 to 0.50 sec time bins (*p<.006), and 1.10-1.20 sec time bins (*p<.03). Lines with shading indicate mean +/− SEM.

To test this, we ran an encoding model on the peri-event calcium data for each rat, for each trial, on the day the animal reached final behavioral criteria in Stage 5 of the DS task. This encoding model was adapted from Parker et al. 2016. Here, a binary of each behavioral epoch was correlated to each time point of the calcium signal, and separate kernels of regression coefficients were acquired for each behavioral epoch (Fig. 4D-F). A time bin analysis was run on each kernel, comparing the average kernel value at each 0.1 second time bin post event onset to a null value, corrected for multiple comparisons. This analysis found that the DS kernel was significantly greater than zero during 0.90-3.0 second time bins post-cue (Fig. 4D; ps<0.05), the PE kernel was significantly greater than zero during 1.40-2.40 second time bins (Fig. 4E; ps<0.05) and the initial lick kernel was significantly increased during 1.30-3.60 second time bins (Fig. 4F; ps<0.05). AUC analysis of the average kernels for each events indicated each kernel is significantly greater than zero (Fig. 4G; DS=1.91+/−0.72, PE=2.10+/− 0.94, lox=1.90+/−0.66; p’s<0.05) Therefore, all three task events (cue onset, reward-seeking, and consumption) are contributing to the VP GABA calcium response.

### VP GABA cue response predicts motivational value of the reward-predictive cue

We next sought to determine whether DS cue-evoked VP GABA calcium activity is correlated with port entry latency on a trial-by-trial basis. Previous work found that the DS-evoked spiking activity of many single units in VP is predictive of the latency it takes the animal to perform the behavior needed to obtain reward, whether that behavior is a lever press or a port entry (Richard et al., 2016, 2018). Because the encoding model described above revealed significant port-entry related increases in calcium activity after, but not before, port entry, for our correlation analysis we only used calcium signals between cue onset and the port entry to examine how cue-elicited VP GABA activity predicted the animal’s latency to enter the port. Latency correlations were calculated for each rat individually in 0.10 second time bins and compared to shuffled controls (Fig. 4H). When we examined the mean correlation coefficients over time, we found that post-DS VP GABA calcium activity is initially positively correlated with port entry latency when compared to shuffled data in the 0-0.50 second time bins (p<.006). This ramping positive correlation is initiated pre-cue onset, during the baseline period. This suggests that baseline activity may predict port entry latency, such that greater baseline VP GABA activity pre-cue leads to a longer post-cue latency to enter the port. However, the 1.10-1.20 second time bins after cue onset are negatively correlated to the latency to enter the port (p<.03). This significant time of negative correlation roughly aligns with kernel time bins in which we observed significant DS-evoked calcium activity (0.90-1.30 second bins). This suggests that VP GABA calcium activity in response to DS cue onset is predictive of port entry latency, such that higher VP GABA calcium activity is associated with shorter port entry latencies, thus greater encoding of the motivational value of the cue.

### VP GABA neurons have a response post reward-predictive cue even in the absence of reward

Because calcium imaging is a proxy for neural activity with slow kinetics and VP neurons have been shown to encode relative reward value, as well as prediction errors (Ottenheimer et al., 2018, 2020a, 2020a), we next aimed to further dissect the nature of the VP GABA neuron calcium activity we observed in response to the port entry and initial lick epochs. First, we introduced a 1 second delay between port entry and reward delivery to further separate the timing of reward-seeking and initial reward consumption. We also included probe trials where the reward-predictive cue was presented but reward delivery did not occur after port entry (Fig. 5A). This allowed us to determine whether calcium activity is elevated post port entry even in the absence of reward, thus allowing us to see if the port entry responses are related to reward consumption or evaluation. Rats were tested for 2 days in sessions in which 50% of DS trials were probe trials, to prevent behavioral extinction. Reward-seeking behavior persisted through both probe days, where the DS port entry probability remained above our response criteria (port entry behavior during at least 70% of DS trials; Fig. 5B). Average VP GABA calcium traces during trials where reward was delivered at a 1 second delay had two distinct peaks, suggesting that the port entry response was delayed somewhat to match the new reward timing (Fig. 5C). Interestingly, this VP GABA response was present even on probe trials, where no reward was delivered. Comparison of the AUC of VP GABA response between the rewarded trials and probe trials revealed no significant difference between the two signals (Fig. 5D; rewarded: 5.38 +/− 0.92, probe: 6.38+/−0.73; F(1,33.99)=0.10, p=0.75; Bayes factor of 4.17 in favor of the null) Therefore, VP GABA neurons still respond to the cue and at the time that reward delivery is expected in the absence of reward. This suggests that VP GABA calcium activity responses after the port entry may be more related to reward anticipation or expectation, than to actual reward consumption or evaluation.

**Figure 5.**
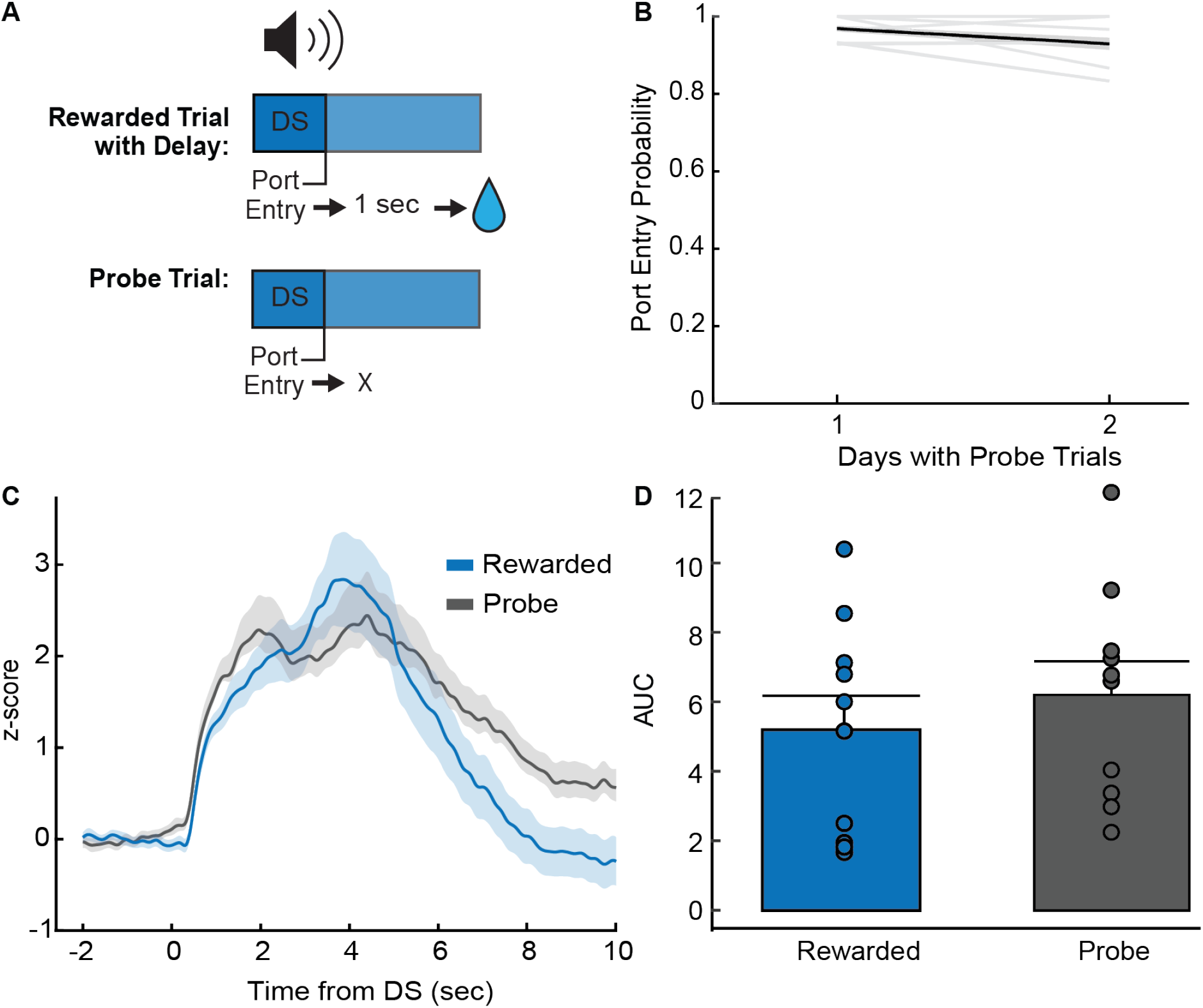
Effects of reward delivery and omission on DS calcium activity. (A) Diagram of delayed reward DS trials and probe DS trials. (B) Average DS port entry probability (n=10 rats) across delayed reward and probe trial sessions (2 days). Behavior remains above criteria of 0.70. (C) Average Z-score peri-DS calcium traces of delayed reward trials and probe trials averaged across animals (n=10) for both days (blue=delayed reward trials, black=probe trials, +/− SEM). (D) Average AUC of post-DS calcium traces (0-5 seconds) for delayed reward (blue) and probe trials (black) across animals (n=10; +/− SEM). We observed no significant difference (sucrose: 5.38 +/− 0.92, probe: 6.38+/−0.73; p=0.78; Bayes factor of 4.17 in favor of the null). Mean + SEM. Dots indicate individual subjects.

### VP GABA neuronal response decreases with extinction of learned behavior

Finally, we examined how consistent omission of reward affected reward-seeking behavior and VP GABA calcium activity. Rats received extinction training for 3 days immediately after probe trial days, where the DS was played but no reward was delivered after port entry. The DS port entry probability of the rats decreased significantly across extinction training days (Fig. 6A; main effect of day: F(2,17)=16.85, p<0.002). VP GABA calcium activity followed a similar pattern, where higher calcium activity in response to DS cue is seen on day 1 of extinction compared to day 2 and 3 of extinction (Fig. 6B). Analysis of the AUC of the traces found a trend toward a significant decrease across extinction days (Fig. 6C; main effect of day effect: F(2,18)=2.77, p=0.10) and planned follow-up comparisons reveal that VP GABA activity on day 2 of extinction training was significantly reduced relative to day 1 (Fig. 6D; p<0.05). This indicates that VP GABA calcium activity is sensitive to extinction learning and changes in the learned value of cues and cue-elicited reward-seeking. Interestingly, both DS-elicited behavior and calcium activity remained above zero, potentially indicating some persistent motivational value of the cue, even after 3 days of extinction.

**Figure 6.**
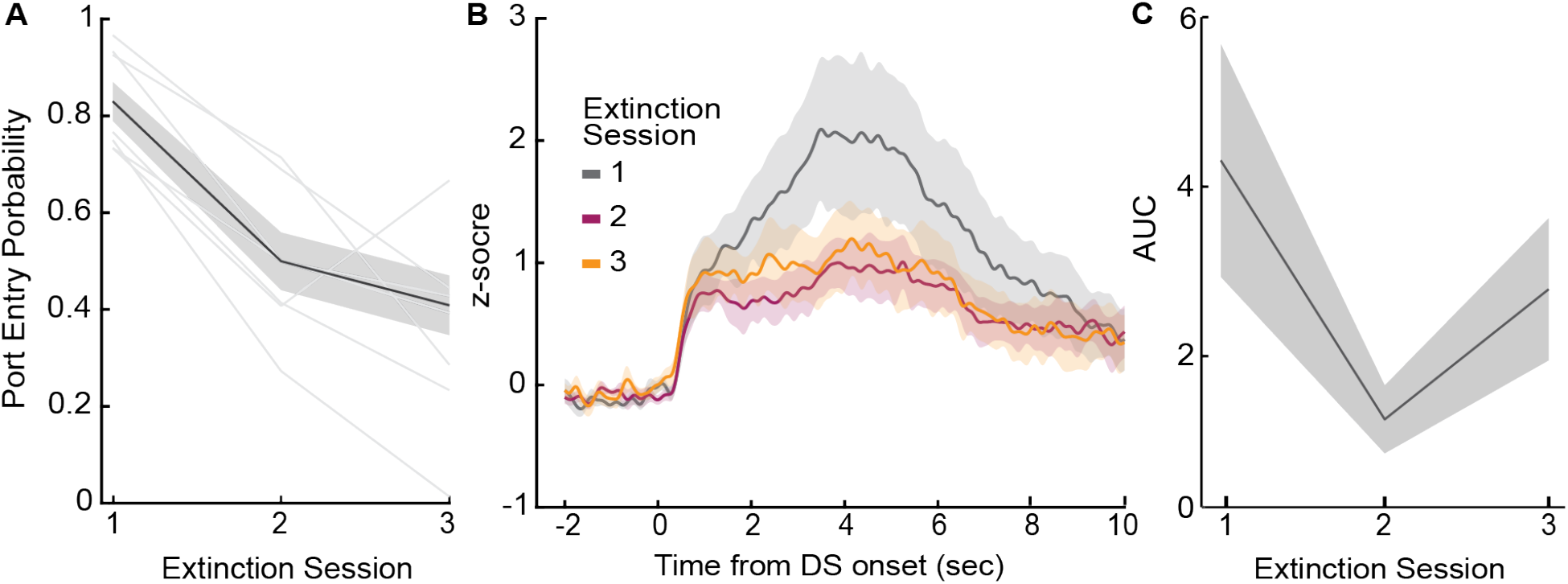
DS calcium activity during extinction learning. (A) Average DS port entry probability across animals (n=8) across extinction sessions (main effect of day: F(2,17)=16.85, p<0.002). (B) Average VP GABA calcium traces across animals (n=8) for each day of training (+/− SEM). (C) Average AUC of post-DS calcium traces (0-5 seconds) across animals (n=8) for each session of extinction training (main effect of day: F(2,18)=2.77, p=0.11). Lines with shading indicate mean +/− SEM. Lines alone indicate individual subjects.

## DISCUSSION

Here we assessed population-level calcium activity in VP GABA neurons while rats learned and performed a cue-elicited reward-seeking task. As rats learned the task, calcium responses developed to the reward-predictive cue but not the neutral cue. An encoding model revealed encoding of the reward-predictive cue, as well as the operant behavior and initial reward consumption. Calcium activity evoked by the reward-predictive cue also predicted the latency of reward-seeking behavior. Calcium responses to the cues and operant behavior persisted even on trials with reward omission. However, when reward was completely omitted from all trials, rats’ behavior, and calcium responses to the reward-predictive cue, decreased. Therefore, VP GABA neurons encode both reward-seeking behavior and the motivational value of reward-predictive cues.

### Ventral pallidal GABA neurons develop calcium responses to cues across learning

Consistent with our overall hypothesis we observed increased calcium activity in VP GABA neurons in response to a cue predicting reward availability, relative to a neutral control cue. Increased firing by these neurons in response to a reward-paired cue has previously been observed in a Pavlovian paradigm (Stephenson-Jones et al., 2020). Specifically, spiking activity of optogenetically-identified VP GABA neurons is elevated in response to a cue predicting water reward in thirsty mice. Additionally, cue-evoked excitatory responses are blunted as mice become satiated on water. These findings suggest that VP GABA neurons may encode the motivational value of cues, but it remained unclear whether this would extend to instrumental reward-seeking in freely moving rats, or to trial-by-trial variations in motivated behavior that are independent of gradual changes in the learned value of the cue due to satiation. Additionally, we wanted to assess the development of these responses across training.

Previous work using *in vivo* electrophysiology to record the activity of VP units in a non-cell-type specific manner found excitatory responses to rewards and reward-related cues in both Pavlovian and instrumental paradigms (Tindell et al., 2004, 2005; Smith et al., 2009, 2011; Ahrens et al., 2016; Richard et al., 2016; Ottenheimer et al., 2018, 2018, 2020a). In a Pavlovian paradigm, the proportion of VP neurons that are excited by a reward-paired cue increases from early to late training as behavioral responses to the cue develop (Tindell et al., 2004). To our knowledge, the development of VP cue responses over time has not previously been reported in an instrumental setting with DS cues. Here DS-evoked calcium activity increases as rats learn the task. Our population level measurements make it unclear whether this finding is due to rate or population level cue encoding, but it is consistent with VP encoding of the learned value of cues in an instrumental task.

Importantly, the observed peri-cue calcium response persisted for multiple seconds after cue onset. The kinetics of even the relatively fast GCaMP6f calcium sensor used here limit our ability to separate out responses to individual behavioral events that occur in close succession relative to *in vivo* electrophysiology (Wei et al., 2020). This is especially true when assessing responses averaged across trials. To more accurately isolate responses to specific behavioral events, and to determine whether peri-DS calcium activity is attributable to the DS specifically, we used linear regression to evaluate how associated each behavioral epoch was with the calcium signal at each timepoint of the task (Parker et al., 2016). With this encoding model, we demonstrate that VP GABA calcium activity encodes the reward-predictive cue. This is consistent with previous work showing increases in VP spiking activity immediately after cue onset in instrumental and Pavlovian paradigms (Tindell et al., 2004; Richard et al., 2016, 2018; Ottenheimer et al., 2018, 2020a; Stephenson-Jones et al., 2020).

### Encoding of reward-seeking latency by VP GABA calcium activity

We also demonstrate that while pre-cue VP GABA calcium activity is positively correlated with trial-by-trial latency, this relationship switches to a negative correlation around the time of DS-evoked calcium activity. We observed this relationship even after excluding calcium activity after port entry. This negative correlation suggests that greater DS-evoked VP GABA calcium activity predicts reward-seeking latency or vigor. Prior work using *in vivo* electrophysiology has also reported a negative correlation between DS-evoked activity and reward-seeking latency (Richard et al., 2016, 2018), as well as a positive relationship with other behavioral measures of cue-evoked motivation, including the speed of the upcoming approach movement and the animal’s proximity to the movement target at cue onset (Lederman et al., 2021). Approximately 20-25% of neurons are reported to have significant negative correlations between post-cue firing and reward-seeking latency; our findings suggest that at least some of these neurons are GABAergic. The positive correlation between pre-cue (and early post-cue) VP GABA activity and trial-by-trial latency reported here was unexpected, as it has not been reported in prior work using electrophysiology, at least on average across VP single units. Whether this correlation is related to our cell-type specific approach, or to calcium activity specifically is an open question.

### Elevation of calcium activity following port entry and during initial reward consumption

The encoding model revealed that VP GABA neuron calcium activity is also evoked by the operant behavior (port entry) required to obtain the sucrose reward, and by initial reward consumption (initial lick). This is consistent with prior work (Richard et al., 2016) showing that VP single units are excited at both the time of an operant response (lever press) and at entry into the reward port. In the Stage 5 version of the DS task, it is difficult to tell whether port entry-evoked calcium activity is related to the action itself, or to reward expectation or evaluation, as the timing of the action (port entry) and reward delivery are confounded. To address this, we added a delay between port entry and reward delivery as well as DS probe trials where no reward was delivered following port entry. When the timing of reward delivery after port entry is delayed, the timing of the calcium activity shifts to match. This suggests that it is not an action related response, but either reward expectation or evaluation. When we separated reward trials from probe trials, we observed very similar calcium responses in neurons no matter the trial type, suggesting that the calcium response is more related to expectation than reward evaluation or hedonic impact.

Finally, the encoding model showed that part of the calcium response was associated with the first lick after rewarded port entry (i.e. the beginning of sucrose consumption). This was expected, as previous work has found that VP GABA neurons respond to rewards (Stephenson-Jones et al., 2020) and have been implicated in consumption of palatable reward in sated animals (Farrell et al., 2021). However, the same VP GABA activity was also observed on probe trials. This differs from previous work showing that a subset of VP neurons are sensitive to reward identity and encode reward value (Ottenheimer et al., 2020a; Stephenson-Jones et al., 2020).

### Caveats and Future Directions

Here, we used population-level calcium imaging to examine encoding of behavioral events. While the time scale of neuronal activation is captured more accurately with electrophysiological recordings, cell-type specific recording is easier to achieve with calcium imaging approaches, as is measurement of activity from neurons with low basal firing rates. We found that VP GABA calcium activity increases in response to a reward-predictive cue and at the time of expected reward consumption. Unexpectedly, we found that the calcium response at the time of expected reward delivery was insensitive to reward omission. This is surprising due to the finding that VP neurons are sensitive to the relative reward value and can encode reward prediction errors (Ottenheimer et al., 2020a). This discrepancy could be due in part to limitations of bulk calcium imaging. Recent work suggests that fiber photometry recordings of striatal calcium activity is more indicative of non-somatic activity (Legaria et al., 2022). Here, we may be capturing dendritic calcium changes, which may not reflect electrophysiological responses that encode relative reward value or reward prediction error signals. Future work could examine soma-specific calcium activity in the DS task using soma-targeted viral expression and/or miniscope imaging (Aharoni and Hoogland, 2019; Chen et al., 2020; Shemesh et al., 2020). Our finding that calcium activity persists even when reward is omitted could also be due to heterogeneity in the response patterns of VP GABA neurons. Importantly, signals encoding relative reward value and reward-prediction error-like signals have just been observed in subsets of VP neurons (Ottenheimer et al., 2018, 2020a, 2020b). VP GABA neurons may differ in their encoding and functional contributions based on more specific cell-type subdivisions or output pathways. For instance, some work points to parvalbumin neurons being a subset of VP GABA neurons involved in cue-evoked behaviors (Prasad et al., 2020), whereas PAS 1-positive VP GABA neurons may contribute to stress and anxiety-related behaviors (Morais-Silva et al., 2022). Additionally, VP GABA neurons have many projection targets that have varied roles in appetitive behavior (Covelo et al., 2014; Mahler et al., 2014; Leung and Balleine, 2015; Root et al., 2015; Prasad and McNally, 2016; Farrell et al., 2021; Vachez et al., 2021; Morais-Silva et al., 2022). Probing a more specific subset of VP GABA cells (cell-type or projection-specific) may reveal greater, or lack of, reward-predictive cue encoding and latency correlations.

## Acknowledgements

This work was supported in part by National Institutes of Health grant R01DA053208 to J.M.R, and a MnDRIVE Graduate Fellowship in Neuromodulation to A.S. Some viral vectors used in this study were generated by the University of Minnesota Viral Vector and Cloning Core.

